# Biogenesis of hepatitis B virus e antigen is driven by translocon-associated protein complex and regulated by conserved cysteines in signal peptide

**DOI:** 10.1101/2021.06.17.448870

**Authors:** Helena Zábranská, Aleš Zábranský, Barbora Lubyová, Jan Hodek, Alena Křenková, Martin Hubálek, Jan Weber, Iva Pichová

**Affiliations:** Institute of Organic Chemistry and Biochemistry of the Czech Academy of Sciences, Flemingovo náměstí 2, 166 10 Prague, Czech Republic

**Author notes:** Address correspondence to Iva Pichova,.

**Keywords:** Hepatitis B virus, HBV precore protein, HBe, signal peptide, cysteine residues, TRAP complex, ER translocation

## Abstract

Hepatitis B virus (HBV) uses e antigen (HBe), which is dispensable for virus infectivity, to modulate host immune responses and achieve viral persistence in human hepatocytes. The HBe precursor (p25) is directed to the endoplasmic reticulum (ER), where cleavage of the signal peptide (sp) gives rise to the first processing product, p22. P22 can be retro-translocated back to the cytosol or enter the secretory pathway and undergo a second cleavage event, resulting in secreted p17 (HBe). Here, we report that translocation of p25 to the ER is promoted by translocon-associated protein complex (TRAP). We found that p25 is not completely translocated into the ER; a fraction of p25 is phosphorylated and remains in the cytoplasm and nucleus. Within the p25 sp sequence, we identified three cysteine residues that control the efficiency of sp cleavage and contribute to proper subcellular distribution of the precore pool.

## Introduction

Hepatitis B is a liver infection caused by the hepatitis B virus (HBV), which can induce both acute and chronic disease and is a major global health problem. According to the World Health Organization, an estimated 257 million people worldwide are infected with HBV. HBV, a member of the *Hepadnaviridae* family, is a small enveloped DNA virus with a genome containing only four open reading frames (C, P, S, and X) that largely overlap and encode multiple proteins using different in-frame start codons. For example, the HBV preC-C gene gives rise to two different products translated from distinct mRNAs – core protein (HBc) and precore protein (HBe). Despite their high sequence similarity, these proteins exhibit different functions and subcellular localizations. HBc is a cytosolic protein with a molecular weight of 21 kDa responsible for the assembly of icosahedral viral particles and pre-genomic RNA encapsidation. On the other hand, the precore precursor, which includes the entire core protein sequence, undergoes a two-step maturation process resulting in production of the extracellular immunomodulatory HBe antigen.

Precore is translated with a 29-amino-acid N-terminal sequence that leads this 25-kDa protein (p25) into the endoplasmic reticulum (ER), where the signal peptide (sp) comprising the first 19 amino acids (the pre sequence) is cleaved off, producing a 22-kDa precore protein (p22) (1). The remaining 10-amino-acid extension at the p22 N-terminus, termed precore propeptide (pro sequence), is not present in the core protein and plays a crucial role in precore folding. An intrasubunit disulfide bond between Cys −7 within the propeptide sequence and Cys 61 changes the dimerization interface, prevents multimerization, and holds the protein in a dimeric state (2–4). The majority of p22 is further processed at its C-terminus by furin protease in the trans-Golgi network, giving rise to mature HBe antigen (p17). The mature antigen is secreted (5–8) and performs an immunomodulatory function in the establishment of persistent infection (9–12).

However, approximately 15–20% of p22 does not enter the secretory pathway and is retrotranslocated back to the cytosol and transported into the nucleus (13–15). The biological function of precore protein in the cytoplasm and nucleus remains poorly understood. P22 can form heterodimers with the core protein and destabilize the viral particle, which may negatively regulate viral infection (16). Conditional expression of precore protein may alter the expression profile of several host genes in transfected hepatocytes (17). Precore also may influence the Rab-7 dependent regulation of HBV secretion (18), promote hepatocellular carcinogenesis by affecting the stability and activity of the p53 tumor suppressor (19), and influence the antiviral signaling of IFN-α (20).

The mechanism by which p22 is distributed to different cellular compartments remains unclear. The SeC61 translocon, together with associated protein complexes, serves as a crossroad for protein translocation into the ER as well as for export via the ER-associated protein degradation (ERAD) pathway [reviewed in (21)]. One member of this machinery, the ER-resident chaperone GRP78/BiP, plays a role in retrotransport of p22 from ER to the cytosol (13). GRP78/BiP participates not only in binding the proteins subjected to ERAD but also in mediating translocon gating (22–24). Another factor that can support protein translocation and SeC61 translocon channel opening in a substrate-specific manner is the translocon-associated protein complex (TRAP), which consists of four subunits (α, β, γ, δ) and supports Sec61 gating for proteins with weak signal sequences (25–28).

Despite recent progress in understanding the intracellular pathways of individual precore forms and growing evidence for their specific roles, many aspects of cytosolic p25 and p22 protein function remain unclear. Furthermore, the mechanism that determines the distribution of precore forms to the secretory or retrotranslocation pathways is not understood. Here, we reveal that conserved Cys residues in the sp sequence are critical for the rate of p25 processing and appear to serve as an auto-regulating factor that influences the intracellular localization of precore. We also describe how the host factor TRAP promotes efficient precore protein ER translocation and prevents mislocalization.

## Results

### A fraction of the cytosolic p25 precursor is phosphorylated

To investigate in detail the process of precore maturation, we performed ^35^S metabolic labeling of cells transiently transfected with an HBe-producing construct (pCEP HBeM1A, resulting protein denoted HBe or precore wt), in which the internal ATG initiation triplet for the core protein (HBc) was mutated (Met/Ala mutation in position 1) (Fig. 1). Huh7 cells were labeled for 30 min, lysed and HBV precore-related proteins were immunoprecipitated with anti-core polyclonal antibody. Although p22 was the predominant species, we also observed a significant portion of unprocessed p25 (around 30% of the total precore signal in pulse samples), indicating that either translocation to the ER or sp cleavage was not 100% efficient. Furthermore, p25 appeared as a double band, suggesting that a portion of it was post-translationally modified (Fig. 2A).

**Fig. 1.**
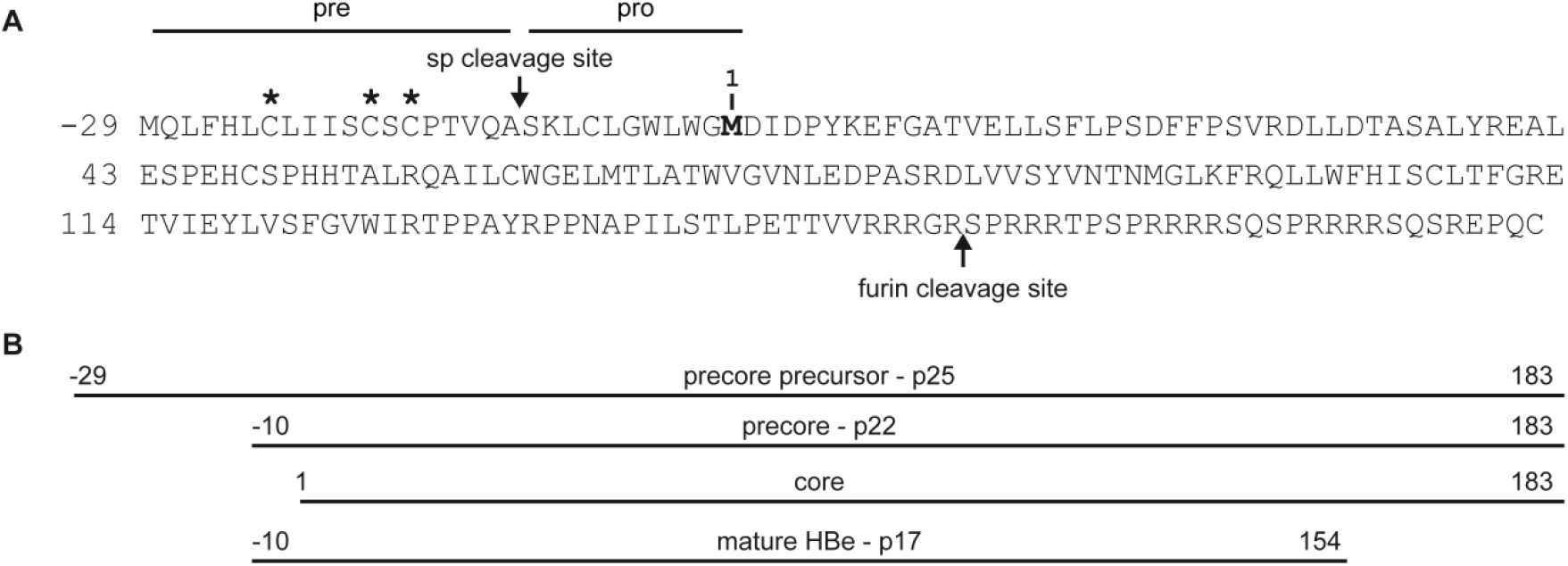
HBV precore precursor and related protein products. (**A**) Sequence of the precore precursor p25. Cysteine residues at positions −23, −18, and −16 are marked by asterisks. The initial methionine of the core protein in position 1 is labeled in bold, and the sites of signal peptide (sp) cleavage and p17 proteolytic maturation are indicated with arrows. (**B**) Schematic representation of individual preC/C gene products.

**Fig. 2.**
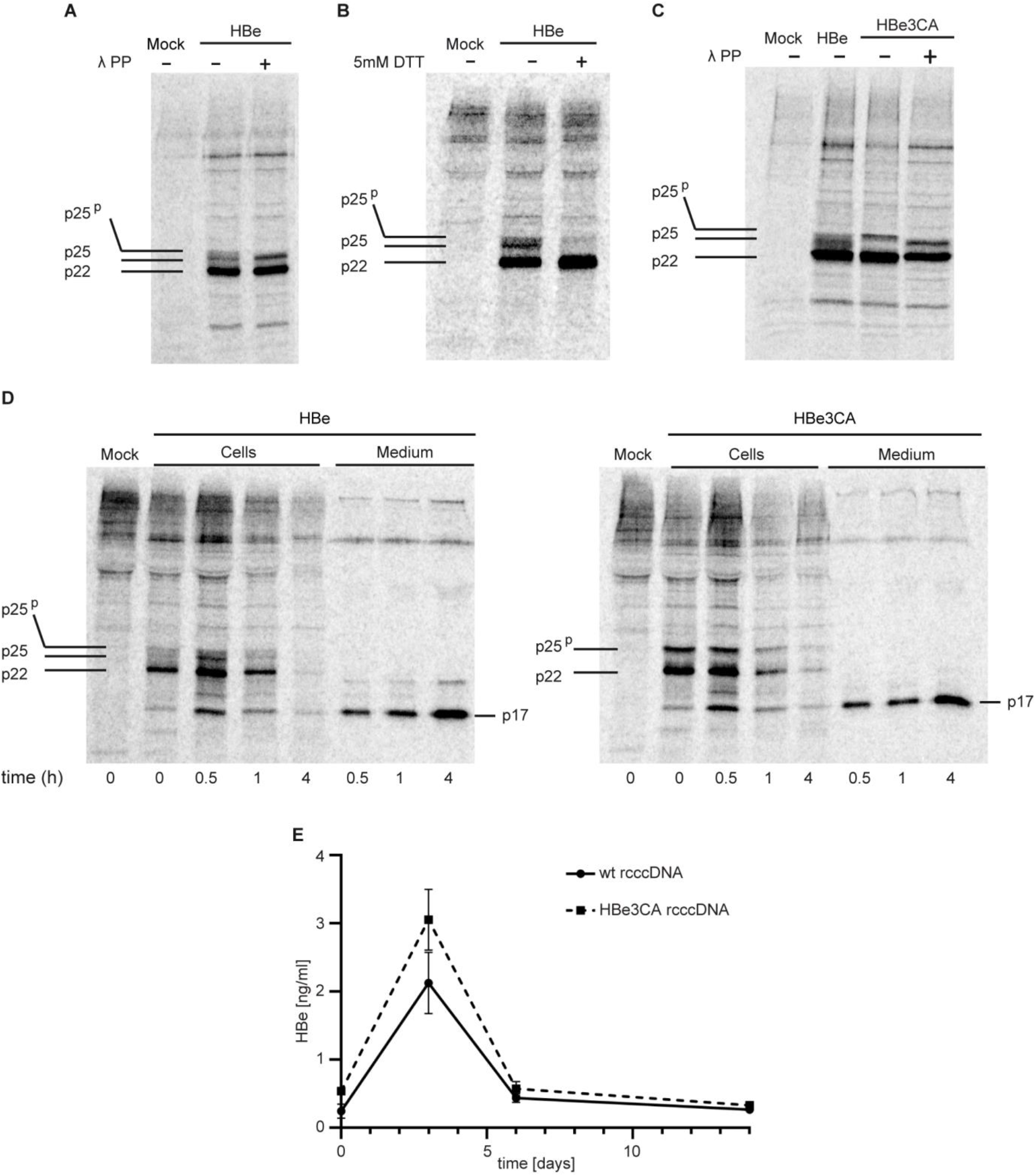
Analysis of p25 protein phosphorylation and effect of mutation of Cys residues in the sp sequence on HBV precore protein processing and virus infectivity. (**A**) Huh7 cells expressing HBV precore precursor were metabolically labeled for 30 min with ^35^S, lysed, and subjected to immunoprecipitation with anti-core antibody. Immunoprecipitated samples were untreated (−) or treated (+) with λ protein phosphatase and separated by SDS-PAGE. The electrophoretic mobility of precore related proteins was analyzed by autoradiography. The migration positions of the HBV precore forms p25^p^, p25, and p22 are indicated. (**B**) Ratios of individual precore protein forms produced in the presence (+) or absence (−) of dithiothreitol (DTT). Huh7 cells expressing the precore precursor were metabolically labeled for 30 min with ^35^S under standard or reducing (5 mM DTT) conditions. HBV precore-derived proteins were immunoprecipitated from the harvested cells with anti-core antibody, separated by SDS-PAGE, and analyzed by autoradiography. (**C**) Electrophoretic mobility comparison of HBe and the HBe3CA mutant. The experiment was performed as for Fig. 2A, (+) phosphorylation of HBe3CA p25 protein was demonstrated by λ-PP treatment. (**D**) Pulse-chase analysis of precore protein processing and secretion. Huh7 cells expressing HBe or HBe3CA proteins were metabolically labeled for 30 min with ^35^S, and then chased for 0.5, 1, and 4 h. At all time points, both cells and media were harvested and subjected to immunoprecipitation with anti-core antibody. Proteins were separated by SDS-PAGE and analyzed by autoradiography. (**E**) Comparison of HBe secretion by cells infected with wild type and mutant precore rcccDNA. HepG2-NTCP cells in 12-well plates were infected in triplicate with wild type rcccDNA (wt rcccDNA) and mutant precore rcccDNA (HBe3CA rcccDNA) with equal MOI of 2000 VGE/cell. Secretion of HBe in the culture supernatants was determined by ELISA at days 0, 3, 6, and 14. Data are plotted as mean ± SEM of three biological replicates.

Because HBV core and p22 are both known to be phosphorylated (29), we investigated the possibility that the upper p25 band (denoted as p25^p^) represents an un-translocated version of p25 that is phosphorylated in the cytoplasm. Treating the immunoprecipitated samples of precore proteins with λ-protein phosphatase (λ-PP) resulted in disappearance of the p25^p^ form, accompanied by enrichment of p25 as visualized by autoradiography (Fig. 2A). These data demonstrate that a significant amount of p25 is not processed by the signal peptidase in the ER and is phosphorylated.

### Reducing conditions or mutation of Cys residues in the sp of p25 stimulates sp cleavage and enhances the rate of p17 biogenesis

As our experiments indicated delayed processing or ineffective translocation of p25, we sought to define the factors that influence the trafficking pathway of the HBV precore precursor. The N-terminal pre sequence of p25, which contains three Cys residues at positions −16, −18, and −23, serves as a sp directing the protein into the ER (see Fig. 1). Although the signal sequence is unusual considering its reduced hydrophobicity, it is conserved among hepadnaviruses (Fig. S1). While Cys −7 within the propeptide sequence is known to stabilize the intrasubunit dimer via a disulfide bond with Cys 61, the role of the three Cys residues located within the sp sequence had remained unclear. We first examined whether the p25 maturation process is influenced by changes in redox conditions. Huh7 cells transfected with pCEP HBeM1A were labeled for 30 min with ^35^S in the presence or absence of 5 mM DTT. The cells were harvested and lysates were subjected to immunoprecipitation with anti-core antibody. In Huh7 cells cultivated without a reducing agent, we observed all intracellular precore forms: p25^p^, p25, and p22. However, in DTT-treated cells, the sp was almost completely removed, and p22 was predominant (Fig. 2B).

Next, we analyzed the contribution of the three Cys residues in the precore precursor sp sequence to p25 processing. We substituted these Cys residues with alanines (construct pCEP HBeM1A C-16A, C-18A,C-23A; the resulting protein is denoted HBe3CA) and transfected this construct into Huh7 cell line, which was labeled for 30 min with ^35^S 24 h after transfection. The autoradiographs of immunoprecipitated proteins (Fig. 2C) indicate that processing of the HBe3CA unphosphorylated precursor was more effective; we observed only phosphorylated p25^p^ and p22 and did not detect any p25. Treatment of these immunoprecipitates with λ-PP resulted in disappearance of the p25^p^ band, further confirming the presence of only the phosphorylated form of p25 in cells transfected with the HBe3CA construct.

To analyze the contribution of the sp Cys residues to the rate of p25 processing and p17 release, we performed pulse/chase experiments with both wt HBe and the HBe3CA mutant. Huh7 cells were isotopically pulse-labeled for 30 min in the presence of ^35^S and chased at different time points. HBV precore-related proteins were immunoprecipitated with anti-core polyclonal antibody (Fig. 2D). In cells producing wt HBe, the amount of p22 decreased significantly after approximately 4 h of chase, which corresponded well with increased extracellular p17 concentration at this time point. In cells producing the HBe3CA mutant, p17 secretion was not impaired, demonstrating that mutation of Cys residues does not interfere with sp function. Moreover, we did not observe the unprocessed unphosphorylated p25 protein in these cells, suggesting more efficient sp cleavage than for the wt.

To explore the function of the N-terminal sequence of precore precursor in the context of the whole virus, mutations of Cys −16, −18, and −23 of p25 were introduced into the HBV recombinant cccDNA minichromosome (3CA rcccDNA). After transfection of wt or 3CA rcccDNAs into HepG2-NTCP cells, the wt and C3A virions were purified from the culture medium. Subsequently, HepG2-NTCP cells were infected with wt and 3CA HBV virions (MOI = 2000 VGE/cell) and the rate of HBV replication was determined by ELISA on days 0, 3, 6, and 14 post-infection. Both wt and 3CA viruses were able to infect HepG2-NTCP, but infection with the 3CA mutant virus yielded higher levels of p17 in the media compared to the wt virus (Fig. 2E). This implies that 3CA mutation may lead to more efficient maturation and secretion of precore protein.

These results indicate a regulatory role for the N-terminal Cys residues in HBV precore protein maturation that is reflected by the rate of p17 secretion.

### Cysteine residues in the sp sequence influence subcellular localization of precore protein

To determine the localization of individual precore protein forms, we performed crude subcellular fractionation of HEK 293T cells, which yielded better separation of individual fractions than Huh7 cells. The cells were transfected with the pCEP HBeM1A or pCEP HBeM1A 3CA construct and isotopically labeled for 30 min with ^35^S. Individual cytosolic, microsomal, and nuclear fractions of the cell lysates were isolated and analyzed by Western blots using antibodies against specific organelle markers (Fig. S2). Precore-related proteins were immunoprecipitated with anti-core polyclonal antibody and visualized by autoradiography. Autoradiographs of HBe samples showed that unphosphorylated p25 and p22 were preferentially localized in microsomes (Fig. 3A), again indicating inefficient and likely post-translational sp cleavage. The main difference between the precore wt and the Cys mutant appeared in the microsomal fraction, in which the HBe3CA samples contained almost no p25, indicating a very fast and effective N-terminal truncation process (confirmed in Huh7 cells; see Fig. S3). To determine whether the inefficient cleavage of wt p25 and its presence in the microsomal fraction is a consequence of a translocation defect and whether the full-length precursor is only attached to the surface of microsomes, we subjected this fraction to proteinase K (PK) cleavage. A sample with 1% Triton 100 added to dissolve all membranes served as a control. After 1 h incubation of microsomes with PK, we still detected undigested wt p25, indicating its membrane shielding and functional translocation (Fig 3B). The phosphorylated p25^p^ form was present in both the cytosolic and nuclear fractions (Fig. 3C), and thus it is tempting to speculate that phosphorylation can stimulate transport of p25 to the nucleus.

**Fig. 3.**
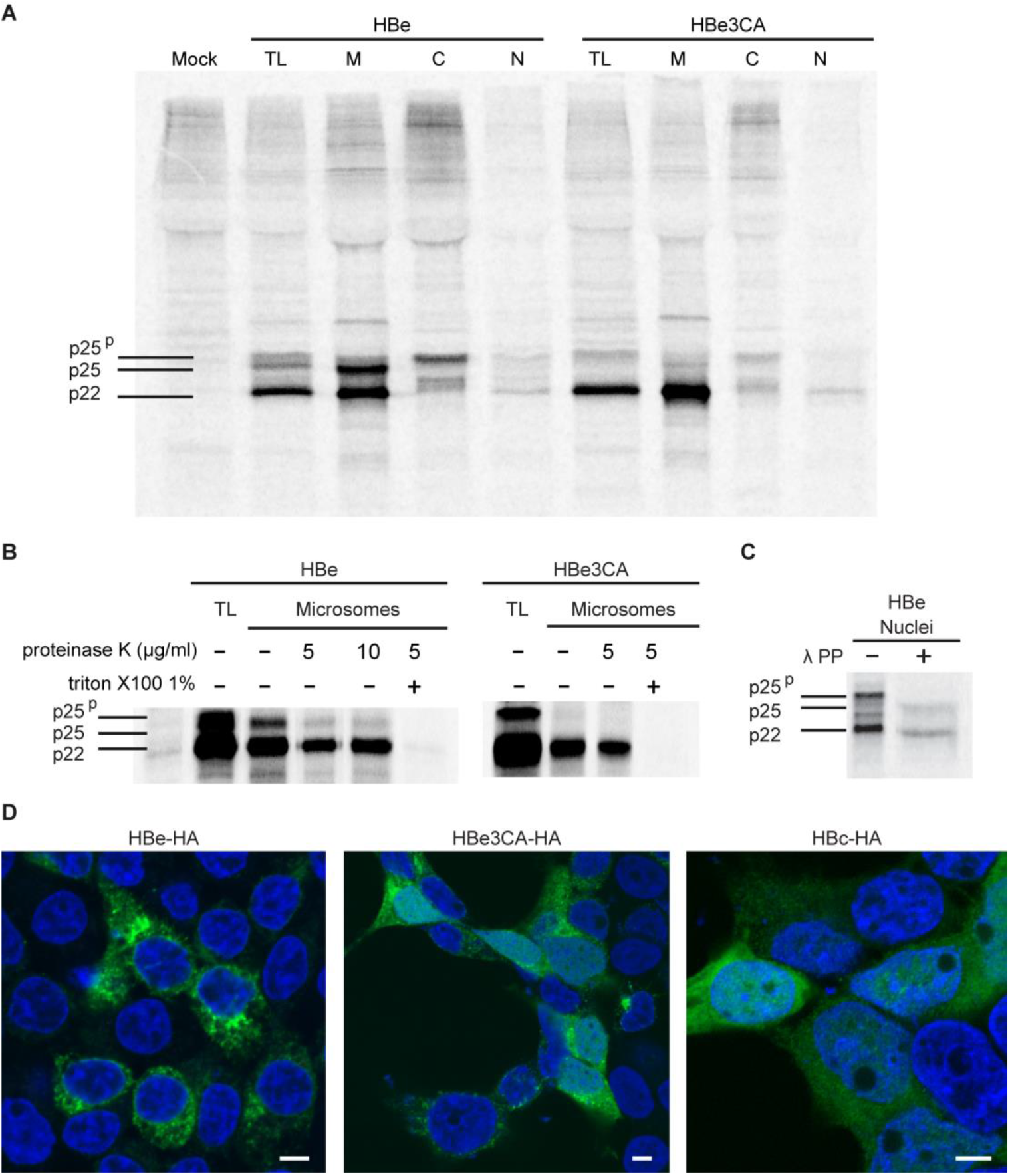
Subcellular localization of wt HBe and HBe3CA-derived precore forms. (**A**) Autoradiograph of ^35^S-labeled proteins immunoprecipitated with anti-core antibody. HEK 293T cells producing HBe or HBe3CA proteins were labeled for 30 min, lysed (total lysate, TL), and subjected to subcellular fractionation. Precore-related proteins from individual fractions representing cytosol (C), nuclei (N), and microsomes (M) were immunoprecipitated, separated by SDS-PAGE, and visualized by autoradiography. (**B**) Immunoprecipitated microsomal fractions from HEK 293T cells were treated with proteinase K and analyzed by autoradiography. (**C**) The immunoprecipitated nuclear fraction from HEK 293T cells was treated with λ-PP separated by SDS-PAGE. The electrophoretic mobility of precore-related proteins untreated (−) or treated (+) with λ-PP was analyzed by autoradiography. (**D**) Representative confocal microscopy images of HA-tagged HBe, HBe3CA, and HBc visualized by FITC-conjugated anti-HA antibody in transfected HEK 293T cells. HBe, HBe3CA, HBc (green) and DAPI (blue). Scale bar represents 5μm.

The subcellular localizations of C-terminally HA-tagged wt HBe, the HBe3CA mutant, and HBc protein (control) were evaluated by immunofluorescence analysis (IFA) using anti-HA antibody (Fig. 3D) in transfected HEK 293T cells. While the wt precore protein was localized exclusively in the cytoplasm and the core protein was distributed between the cytosol and nucleus with predominant nuclear localization, the HBe3CA mutant displayed a mixed phenotype. We observed cells with a cytoplasmic phenotype resembling that of wt, as well as cells exhibiting both a nuclear and cytoplasmic distribution pattern. We assume that the increased level of HBe3CA mutant in the nucleus is related to a higher intracellular level of p22 resulting from faster sp processing, which could contribute to the massive retro-translocation from the ER.

To examine whether the HBV precore signal peptide can work independently of the rest of the protein sequence, we prepared constructs of enhanced green fluorescent protein (EGFP, vector pEGFP N1) N-terminally attached to the leader sequences and propeptide sequences from either wt p25 or the 3CA mutant [prepro sequence, amino acids (−1) – (−29), resulting constructs pCMV preproHBe-EGFP and pCMV preproHBe3CA-EGFP]. As a control, we prepared a similar construct with a leader sequence plus four additional amino acids from the mature part of the human protein disulfide isomerase PDIA1, an abundant ER resident chaperone (attached sequence MLRRALLCLAVAALVRADAPE, construct pCMV spPDI-EGFP). The subcellular localization of these constructs in transfected HEK 293T cells was evaluated by confocal microscopy (Fig. S4). The localization pattern of wt EGFP appeared very dispersive, as the protein migrates by free diffusion from the cytosol to the nucleus. The phenotypes of the preproHBe-EGFP and preproHBe3CA-EGFP constructs were similar to that of EGFP, implying that HBV precore sp (either wt or mutated) is not sufficient for successful ER translocation and that the cooperation of downstream sequences in this process is likely necessary. In contrast, the pattern of EGFP fused with the functional PDIA1 leader sequence exhibited cytoplasmic localization in about 50% of transfected cells. Apparently the efficiency of HBV precore signal sequence alone is not comparable with that of the common leader signal from an ER-resident protein.

Taken together, these data indicate that precore protein translocation is a controlled process, in which the delay in p25 N-terminal cleavage and thus regulation of the p22 level prevent its mislocalization within cells. Cys residues in the sp and the weak ability of the leader sequence to mediate translocon gating are likely key factors in this sequential HBe maturation.

### HBV precore precursor interacts with TRAP

To identify host proteins potentially involved in translocation of the HBV precore protein, we performed HBe interactome analysis using an LC-MS/MS-based proteomics approach. To block HBe secretion and thereby enhance its detection in cell lysates, we added Brefeldin A, which inhibits protein transport from the ER to the Golgi complex. HepG2-NTCP cells transfected with a plasmid encoding C-terminally HA-tagged HBe or HBe3CA were cultivated for 36 h, treated with Brefeldin A for 4 h, and harvested. Cell lysates were subjected to immunoprecipitation using anti-HA magnetic beads, and recovered proteins were analyzed by LC-MS/MS for identification and label-free quantification. Triplicates of HBe (or HBe3CA) and control cell lines allowed application of a Student’s *t*-test to statistically determine the proteins specifically enriched in HBe positive samples. The result of the analysis is shown on the Volcano plot (Fig. S5). Among the proteins enriched in HBe (or HBe3CA) containing samples we identified the previously described precore interacting partners C1qBP, GRP78/BiP (13,30), and protein kinase SRPK1, which mediates HBV core phosphorylation (31) (Tab. 1). In both HBe-HA and HBe3CA-HA samples, we also repeatedly observed peptides derived from the Sec61 translocon complex and subunits (α, β, δ) of TRAP, an accessory component that triggers Sec61 channel opening in a substrate-specific manner (Tab. 1) (27). No significant differences were observed between wt precore and the 3CA mutant with regard to detected co-immunoprecipitated proteins.

**Table 1:** HBe associated proteins identified by shotgun LC-MS/MS analysis of HBe-HA immunoprecipitates from HepG2 cells.

| Protein | HBe<br>Peptides<br>Exp.1/2/3 | HBe<br>PSMs<br>Exp.1/2/3 | CTRL<br>Peptides<br>Exp.1/2/3 | CTRL<br>PSMs<br>Exp.1/2/3 |
| --- | --- | --- | --- | --- |
| **HBe-HA** | **17/13/16** | **45/35/45** | **0/0/0** | **0/0/0** |
| C1qBP | 9/8/9 | 30/23/36 | 0/0/1 | 0/0/1 |
| GRP78/BiP | 6/5/7 | 10/9/11 | 4/3/0 | 6/4/0 |
| SRPK1 | 3/4/5 | 4/5/7 | 0/0/0 | 0/0/0 |
| Sec61 alpha-1 | 2/1/2 | 5//2/3 | 2/0/1 | 2/0/1 |
| Sec61 alpha-2 | 1/0/1 | 3/0/1 | 0/0/0 | 0/0/0 |
| TRAP alpha | 2/1/1 | 5//4/4 | 0/1/1 | 0/2/2 |
| TRAP beta | 2/1/1 | 3/2/1 | 1/0/0 | 1/0/0 |
| TRAP delta | 1/1/0 | 1/2/0 | 0/0/0 | 0/0/0 |

(HBe, HBe-HA sample; CTRL, negative control (no HBe) sample; peptides, the number of detected peptide sequences unique to a protein; PSM, the total number of identified peptide spectra matched for the protein)

To evaluate potential interactions between the HBV precore protein and TRAP complex, we co-transfected HEK 293T cells with the HA-tagged HBe construct and individual C-myc-tagged TRAP subunits α, β, γ, and δ and performed pull-down experiments using anti-HA magnetic beads followed by Western blot evaluation using anti-c-myc antibody (Fig. 4A). We detected a significant signal corresponding to all four individual TRAP subunits after co-immunoprecipitation with the HBe construct, indicating mutual interaction between HBV precore and TRAP. In control samples transfected with mock DNA and individual TRAP subunits, we observed a non-specific interaction between TRAP β and the magnetic beads; the other three subunits did not display any nonspecific background. The repeatedly confirmed association between the HBV precore protein and TRAP complex subunits strongly implies involvement of TRAP in the translocon gating of p25 protein.

**Fig 4.**
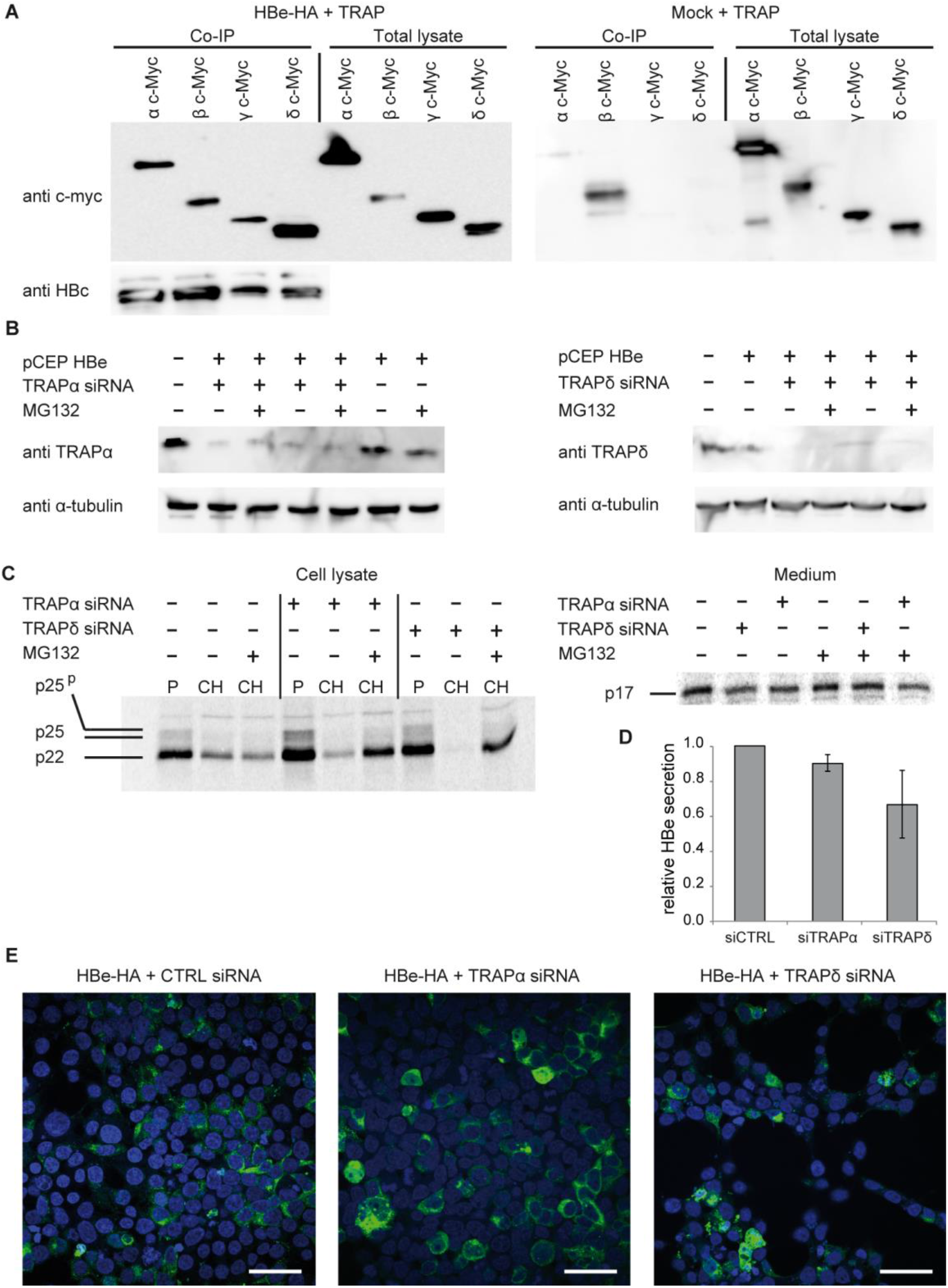
Co-immunoprecipitation of HBV precore protein with individual TRAP subunits and TRAP depletion effect on precore stability and translocation. (**A**) HEK 293T cells were co-transfected with HA-tagged HBe producing construct (or mock DNA) and plasmids expressing individual c-myc tagged TRAP subunits. Precore protein was immunoprecipitated with anti-HA magnetic beads and samples were analyzed by Western blot using anti c-myc antibody to detect interacting TRAP proteins. (**B**) HEK 293T cells were co-transfected with HBe producing construct and siRNAs targeting either TRAP α or δ genes. Silencing effect was evaluated by Western blot analysis. (**C**) Knock down of TRAP decreases efficiency of HBe secretion. 48 h post-transfection, TRAP-silenced cells were metabolically labeled for 30 min with ^35^S (P) and chased for 4 h (CH) with or without addition of the proteasome inhibitor MG132. Cells and media were harvested and subjected to immunoprecipitation with anti-core antibody. Proteins were separated by SDS-PAGE and analyzed by autoradiography. (**D**) The signal intensity of p17 in the medium was quantified and shown relative to the unsilenced sample. The bars represent averages from three independent experiments, and error bars indicate the standard deviation. (**E**) Representative confocal microscopy images of HA-tagged HBe or HBe3CA in HEK 293T cells depleted for TRAP δ subunit visualized by FITC-conjugated anti-HA antibody. HBe and HBe3CA are shown in green, and DAPI in blue. Scale bar represents 40μm.

### TRAP complex cooperates in the translocation and sequential maturation of HBV precore protein

To investigate whether the TRAP complex is involved in precore protein translocation into the ER, siRNA-mediated knockdown of either the α or δ subunit was performed in HEK 293T cells. Cells were co-transfected with plasmid producing HBe and siRNA targeting one TRAP subunit (α or δ). Metabolic labeling followed by a pulse-chase experiment (30 min pulse, 3 h chase) was performed 48 h after transfection. The efficiency of silencing was evaluated by Western blot using monoclonal antibodies that recognize the hTRAP α or δ subunit (Fig. 4B). Cell lysates and collected medium were immunoprecipitated using polyclonal anti-HBV core antibody. The remarkable effect of TRAP depletion became evident at the intracellular precore protein level. In cells producing wt precore protein with silencing of either the TRAP α or δ subunits, the signals of p22 and p25 significantly decreased compared to nondepleted cells and were restored after addition of the proteasome inhibitor MG132. The effect was stronger upon TRAP δ knock down (Fig. 4C). In the medium of cells overexpressing HBV precore with TRAP silencing, we observed slightly lower levels of mature p17 secretion, again especially in samples with TRAP δ depletion (Fig. 4D). Our results indicate that silencing of individual TRAP subunits promotes degradation of HBe wt, presumably due to ineffective translocation.

Next, we analyzed the effect of TRAP depletion on the subcellular distribution of the precore protein. HEK 293T cells were co-transfected with the HBe-HA producing construct and either TRAP α- or δ-targeting siRNA and examined by confocal microscopy after immunofluorescence staining using anti-HA antibody conjugated with FITC (Fig. 4E). In both TRAP-silenced samples, the typical ER localization pattern of precore protein was disrupted in fraction of cells. The protein appeared to be distributed between the cytosol and the nucleus, indicating a certain degree of malfunction in the translocation process.

It is evident that the leader sequence of the HBe p25 precursor is weak to mediate an autonomous translocation into ER and that the assistance of the TRAP complex in conducting channel gating is indispensable for proper HBV precore subcellular localization and p17 biogenesis.

### Silencing of individual TRAP complex subunits in HBV-infected cells reduces the extracellular level of HBe

To determine whether TRAP complex mediates co-translational translocation of HBe precursor in HBV-infected hepatocytes, we examined the consequences of siRNA-mediated knockdown of TRAP subunits in HepG2-NTCP cells. SiRNA-treated HBV-infected hepatocytes were cultured for 5 days before HBe secretion was determined by ELISA. Whereas non-targeting siRNA had no effect on HBe secretion following knockdown, siRNAs targeting the α, β, and δ TRAP subunits reduced the extracellular level of HBe (Fig. 5A). In contrast, the level of HBs antigen secreted from TRAP-silenced cells remained comparable to that from non-silenced cells (Fig. S6A). These results indicate that the stability of TRAP is dependent on the presence of all its components and that the efficiency of HBe secretion is reduced upon downregulation of TRAP in HBV-infected cells.

**Fig. 5.**
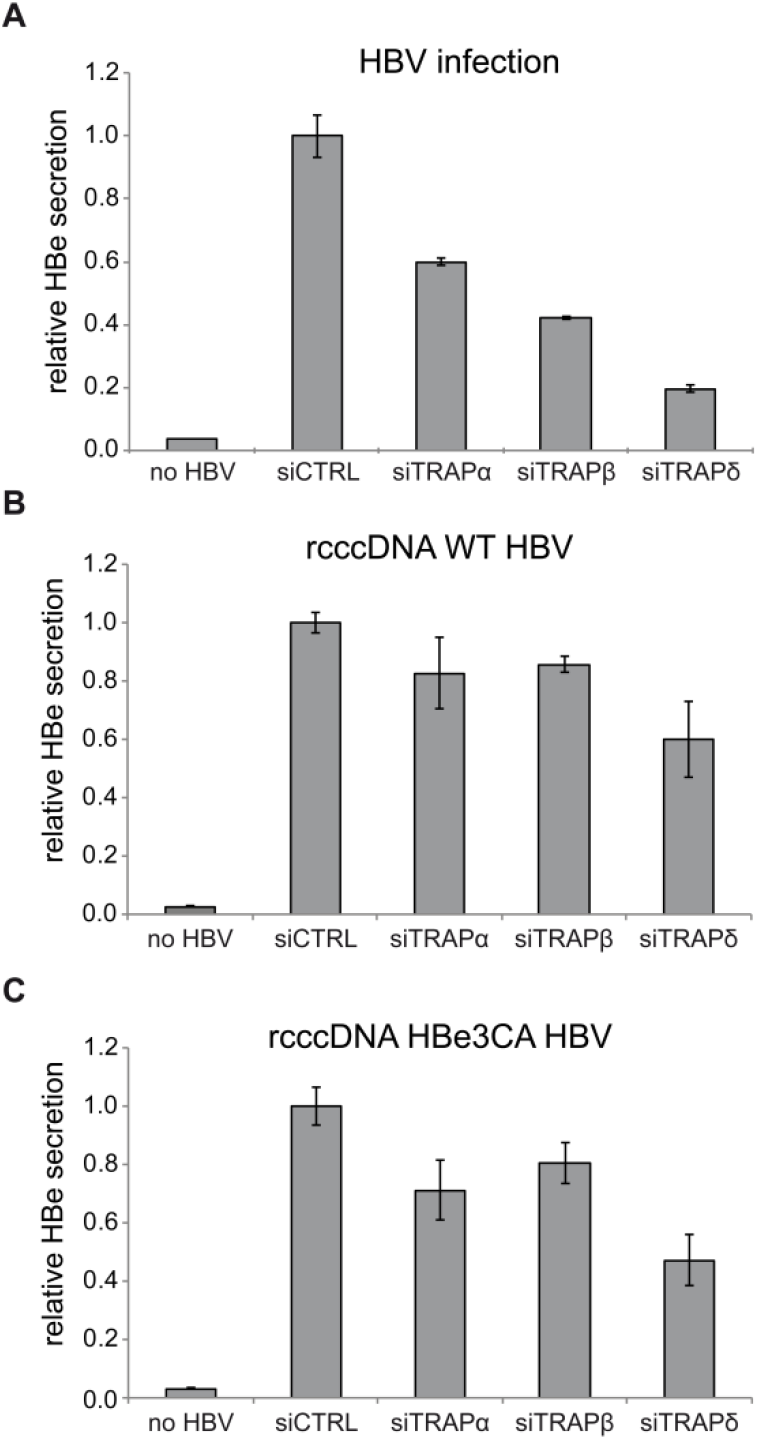
The depletion of TRAP subunits downregulates the extracellular level of HBe in both HBV-infected and wt HBV or HBe3CA rcccDNA transfected hepatocytes. (**A**) Biological triplicates of HepG2-NTCP cells were first transfected with siRNA oligonucleotides for 24 h, then infected with HBV at a MOI of 1,500 VGE/cell for another 96 h before cells and media were harvested. Secreted HBe in media was quantified by ELISA. TRAP subunit knockdown affects HBe secretion for both wt (**B**) and mutant HBe3CA (**C**) HBV. HepG2-NTCP cells were simultaneously transfected with rcccDNA plasmid coding for either WT or HBe3CA virus and siRNAs targeting individual TRAP subunits. Secreted HBe levels in media were quantified by ELISA after 5 days of cultivation.

To exclude the possible cytotoxic effect of individual knockdowns, we analyzed the general effect of TRAP silencing on cell viability. HepG2-NTCP cells were transfected with individual siRNAs, and we included 5% DMSO as a positive control. After 5 days of incubation, cell viability was determined by XTT assay in biological triplicates (Fig. S6B). Other than slightly reduced metabolic viability in cells treated with TRAPδ siRNA, no significant changes were observed. The efficiency of silencing was monitored by RT-qPCR 3 days post-transfection (Fig. S6C).

To determine possible differences between HBe and HBe3CA regarding TRAP depletion in HBV-producing cells, we co-transfected HepG2-NTCP cells with HBV rcccDNA (containing either HBe or HBe3CA genes) and siRNAs targeting individual TRAP subunits. The resulting level of mature p17 secretion was evaluated by ELISA after 5 days. In this experimental setup, we observed reduced HBe levels in the media of TRAP-depleted cells for both wt rcccDNA (Fig. 5B) and HBe3CA rcccDNA (Fig. 5C).

## Discussion

By closely examining the maturation of the HBe precursor p25, we demonstrate here that HBV precore protein is present in cells in different forms and is localized to different subcellular compartments. We identified the unprocessed p25 precursor in microsomes and its phosphorylated form (p25^p^) in the cytosol and nucleus. We found that the three Cys residues in positions −16, −18, and −23 play a coordinated regulatory role that contributes to proper p25 processing and localization. Furthermore, our results illustrate the importance of the host TRAP complex for HBV precore precursor ER translocation.

Phosphorylation and dephosphorylation of HBV core protein plays a crucial role in assembly, disassembly, and nuclear localization of capsids (32–36). Nuclear localization of HBc has been associated with two co-dependent nuclear localization signals within the arginine-rich domain (37,38), which by analogy are also present at the C-terminus of precore (p25 and p22). Nuclear localization of phosphorylated p22 has been described (29,39); our results revealed that the phosphorylated version of p25 also is localized in the nuclear fraction of mammalian cells, although its role in the nucleus has yet to be elucidated. Locarnini and colleagues observed that precore protein induces repression of many genes in transfected hepatocytes and behaves like a tumor-suppressing protein with anti-apoptotic activity (17). However, this observation was presented as a general effect of precore expression in cells, and no particular activity was attributed to individual forms of precore protein.

Factors involved in retaining a significant portion of p25 in the cytoplasm or causing retro-translocation of p25 from the ER membrane back to the cytosol also remain poorly understood. Our findings show that addition of a reducing agent (DTT) into cell culture media or mutations of three Cys residues in the sp sequence accelerate p25 processing and lead to almost complete conversion of p25 to p22. Four cysteines in the N-terminal part of p25 are conserved in all hepadnaviruses, and these sequences are predicted to be potential zinc fingers. Their function in the viral life cycle has not been established. Wasenauer *et al*. reported that these Cys residues are dispensable for mature HBe production (40). In agreement with their data, we found that replacement of three cysteines in the sp sequence with alanines did not abolish precore maturation. The sp function of the HBe3CA construct remained unchanged, and the mutated protein was translocated into the ER. Single-round infectivity assays revealed a slightly higher signal of mature p17 secreted from HepG2-NTCP cells infected with HBe3CA HBV particles compared to cells infected with the wt HBV genome. Interestingly, confocal microscopy experiments showed that Cys residues in the sp are responsible for the proper precore localization pattern and prevent spreading of precore across the cell in a core-like phenotype. It appears that HBV needs to tightly regulate the precore precursor maturation process to avoid unfavorable p22 localization and its eventual interference with core functions. Additionally, the distribution of the dual p25 population (phosphorylated in cytosol and nucleus, nonphosphorylated in microsomes) must be under some control mechanism. Several ER-resident proteins, such as calreticulin, have been found to have another independent role in cytosol, and their segregation between compartments is influenced by the weak ability of their signal sequences to mediate translocation (41,42). Our results showed that p25 processing and localization is regulated by the conserved Cys residues within the sp. In the co-translational pathway, the translocation process is initiated during protein synthesis when the sp is recognized by SRP. As long as this sequence is sufficiently hydrophobic and helical (so-called “strong”), it can facilitate translocon opening. The tertiary folding is delayed until the protein exits the channel and enters the ER lumen [reviewed in (43)]. Domains folded prematurely in the cytosol inhibit translocation (44,45). Gating of proteins with “weak” signal sequences is postponed, and their interactions with additional Sec61 accessory proteins such as TRAM, TRAP, Sec62/63, or BiP are needed [reviewed in (46)]. The composition of the sp of HBV precore is unusual and does not contain a segment of hydrophobic amino acids characteristic of typical leader sequences. Apparently, this signal is not strong enough to open the translocon channel, and a host chaperone is employed to mediate successful gating or to stabilize the binary Sec61–p25 complex. This hypothesis is in agreement with the observation that p25 displays impaired sp cleavage in a non-mammalian model, possibly caused by a lack of appropriate cellular partners (47). As our data suggest that p25 sp does not work independently of the rest of the precore protein, additional downstream sequences are very likely necessary for successful translocation, and we assume that they cooperate with transmembrane or lumenal proteins.

Our work establishes TRAP complex as a novel interacting partner participating in p25 translocation. After siRNA-mediated knockdown of TRAP complex, translocation of wt precore protein precursor is disrupted, and the protein is directed to proteasomal degradation. In HBV-infected hepatocytes, this is reflected by significantly decreased production of secreted p17. Several signal sequences, including those of the prion protein and preproinsulin, require TRAP complex for successful conducting channel opening (48,49). In addition to low hydrophobicity, a high GP content has been proposed as a common feature for most such signal sequences (27). Interestingly, the sp of HBV p25 fulfils only the condition of a reduced number of hydrophobic residues. Instead of a GP-rich patch, its primary structure includes three conserved cysteines, which seem to be involved in autoregulation of HBe biogenesis. Their precise role and possible participation in tertiary structure still need to be elucidated, but evidently, they are not indispensable for TRAP interaction and facilitation of translocon gating. Instead, they are more likely responsible for moderation of the translocation process and thus its regulation.

In summary, we present evidence that the three conserved Cys residues in the sp of the HBV precore precursor p25 serve as an auto-regulatory factor controlling proper progression of the precore translocation process and therefore preventing mislocalization of precore. We also identified TRAP complex as a host factor required for successful translocon gating of p25 and for support of mature p17 biogenesis.

## Experimental procedures

### DNA constructs

HBV precore and core coding sequences were derived from the HBV genome-containing plasmid pHBV4.1 obtained from Dr. Huiling Yang (Gilead Sciences, Foster City, CA, USA). The HBV core protein coding region was amplified using the forward primer 5’-ATCATAAGCTTACCATGGACATCGACCCTTATAAAG-3’ and reverse primer 5’-TAGATGGTACCCTAACATTGAGGTTCCCGAG-3’ and subcloned into the pcDNA3.1 vector (Clontech) via *Hind*III and *Kpn*I restriction sites, giving rise to construct pcDNA3 HBc. The precore precursor gene was amplified using the forward primer 5’-ATCTAAAGCTTACCATGCAACTTTTTCACCTCT-3’ and reverse primer 5’-TAGATGGATCCCTAACATTGAGGTTCCCGAG-3’ and introduced into the pCEP vector (Invitrogen) via *Hind*III and *Bam*HI restriction sites. To remove the initiating ATG codon for the HBV core protein, a M1A mutation was introduced by site-directed Pfu mutagenesis using mutagenic primers 5’-TGGCTTTGGGGCGCCGACATTGACC-3’ and 5’-GGTCAATGTCGGCGCCCCAAAGCCA-3’, giving rise to the construct pCEP HBeM1A.

The C-terminal HA tag was introduced into the pCEPHBeM1A construct using the reverse primer 5’-AGCTCGGATCCTCAAGCGTAGTCCGGGACGTCGTAAGGGTAACATTGAGGTTCCCGA G-3’. The following forward primer was used for mutation of three cysteines at positions −23, −18 and −16 of the precore signal peptide sequence to give rise to construct pCEP HBeM1A3CA: 5’-ATCTAAAGCTTACCATGCAACTTTTTCACCTCGCCCTAATCATCTCTGCTTCAGCTCCTACTGTTCAAGC-3’.

The minicircle-producing plasmid for wt HBV recombinant cccDNA (*wt* rcccDNA) was kindly provided by Dr. Ping Chen (Shenzhen Institutes of Advanced Technology, Chinese Academy of Sciences, Shenzhen, China) (50). The rcccDNA plasmid designated as C3A rcccDNA, which carries a triple C-to-A mutation in the pre-core ORF at positions −16, −18, and −23, was generated by site-directed mutagenesis (QuikChange II XL Site-Directed Mutagenesis Kit, Agilent Technologies) with three subsequent rounds of PCR using the following primers: C1A-F, 5’-GCGATGCAACTTTTTCACCTCGCCCTAATCATCTCTTGTTCATG-3’; C1A-R, 5’-CATGAACAAGAGATGATTAGGGCGAGGTGAAAAAGTTGCATCGC-3’; C2A-F, 5’-CCATGCAACTTTTTCACCTCGCTCTAATCATCTCTGCTTCAG-3’; C2A-R, 5’-CTGAAGCAGAGATGATTAGAGCGAGGTGAAAAAGTTGCATGG-3’; C3A-F, 5’-CAACTTTTTCACCTCGCCCTAATCATCTCTGCTTCAGCTCCTACTGTTCAAGCC-3’; C3A-R, 5’-GGCTTGAACAGTAGGAGCTGAAGCAGAGATGATTAGGGCGAGGTGAAAAAGTTG-3’.

Constructs of individual c-myc tagged human TRAP subunits (α, β, δ) under the CMV promotor were purchased from OriGene Technologies (#RC202408, #RC213580, #RC201079, #C201593).

### rcccDNA production and purification

The wt and C3A mutant rcccDNA plasmids were transformed into *E coli*. strain ZYCY10P3S2T, and the rcccDNA was isolated using the MC-Easy minicircle production kit (System Biosciences) according to the manufacturer’s recommendations.

### siRNA

SiRNAs targeting individual hTRAP subunits (α, β, γ, δ) were purchased from Santa Cruz Biotechnology (SCBT, t# sc-63153, #sc-63147, #sc-63148, #sc-63149).

### Antibodies

Rabbit polyclonal anti-HBV core protein antiserum was obtained after immunization of three animals with 1.4 mg/ml purified denatured (boiling in 1% SDS, 0.1 M DTT) HBV core protein produced in *E. coli* (Moravian Biotech).

Mouse monoclonal antibodies against individual TRAP subunits were purchased from SCBT (#sc-373916, #sc-517428, #sc-376706).

### Cells

HEK 293T (human embryonic kidney cell line, ATCC) and Huh7 cells (differentiated hepatocyte-derived carcinoma cell line, Japanese Collection of Research Bioresources Cell Bank) were cultured in Dulbecco’s Modified Eagle Medium (DMEM) supplemented with 10% fetal bovine serum (FBS, VWR) and an antibiotic mixture (penicillin/streptomycin (PenStrep), Sigma-Aldrich) at 37 °C in a 5% CO_2_ atmosphere.

HepG2-NTCP, a HepG2 human liver cancer cell line stably transfected with the human HBV entry receptor (sodium taurocholate co-transporting polypeptide, NTCP), was obtained from Dr. Stephan Urban (Heidelberg University Hospital, Heidelberg, Germany). The cells were cultured in DMEM supplemented with 10% FBS, the antibiotic mixture (PenStrep), and puromycin (0.05 mg/mL, Sigma-Aldrich) at 37 °C in a 5% CO_2_ atmosphere.

### Cell proliferation assay

Labeling reagents XTT (sodium 3’-[1-(phenylaminocarbonyl)- 3,4-tetrazolium]-bis (4-methoxy6-nitro) benzene sulfonic acid hydrate) and PMS (N-methyl dibenzopyrazine methyl sulfate) were purchased from Sigma-Aldrich. Assay was performed according to protocol in Cell proliferation kit II (XTT) (Roche, #11 465 015 001).

### Transfection

According to the respective manufacturer’s instructions, plasmid DNA was introduced into HepG2-NTCP cells using Lipofectamine™ 3000 transfection reagent (ThermoFisher Scientific), Huh7 cells were transfected with GenJet™ (SignaGen Laboratories), and HEK 293T cells were transfected with a 1:3 ratio of DNA:polyethylenimine (PEI, 25 kDa linear, Sigma-Aldrich). Transfection of siRNA was performed using X-tremeGENE siRNA Transfection Reagent (Roche) for HEK 293T cells and Lipofectamine RNAiMAX Transfection Reagent (ThermoFisher Scientific) for HepG2-NTCP cells, according to the manufacturers’ protocols.

### RT-qPCR

Intracellular levels of mRNA for TRAP complex subunits were estimated by RT-qPCR using Luna Universal One-Step RT-qPCR Kit (NEB) following total RNA isolation with RNeasy Mini Kit (Qiagen). Primers targeting TRAPα, β, and δ mRNAs were purchased from SCBT (#sc-63153-PR, #sc-63147-PR, #sc-63148-PR). Samples were analyzed as technical duplicates.

### Metabolic labeling and pulse/chase analysis

Confluent Huh7 or HEK 293T cells grown on 60 mm dishes were starved 18 h post-transfection with DMEM lacking cysteine and methionine for 15 min, pulsed for 30 min with 10 μCi [^35^S]-labeling mix (Trans^35^S-Label™ MP Biomedicals) and subsequently chased with complete DMEM medium for the desired time periods. Harvested cells were lysed in lysis buffer A (1% Triton X100, 50 mM NaCl, 1% deoxycholate, 25 mM Tris, pH 8) containing 1 mM PMSF and Complete™ EDTA free protease inhibitor cocktail (Roche). Nuclei were removed by centrifugation for 1 min at 6,000 rpm, and immunoprecipitation of the precleared lysate was performed using polyclonal anti-core antibody and protein A sepharose beads (Invitrogen). Immunoprecipitated complexes were washed twice in buffer B (1% Triton X100, 50 mM NaCl, 1% deoxycholate, 0.1% SDS, 25 mM Tris, pH 8) and once in 20 mM Tris, pH 8. Sepharose beads were resuspended in SDS-PAGE loading buffer and boiled for 5 min. Proteins were separated on 15% SDS-PAGE and subjected to phosphorimager analysis. Signal intensity was evaluated using QuantImage software (GE Healthcare).

### Immunofluorescence analysis

Transiently transfected HEK 293T cells were grown on 22-mm glass coverslips for 24 h, briefly washed with phosphate-buffered saline (PBS), and fixed with 4% paraformaldehyde in PBS for 30 min at room temperature. Fixed cells were washed with PBS and permeabilized with 0.2% Triton X-100 in PBS for 30 min. Permeabilized cells were immunostained in PBS, 0.2% Triton X-100, 10% FBS for 1 h using anti-HA antibody conjugated with FITC (Sigma-Aldrich, #H7411). The cells were washed three times for 10 min with 0.2% Triton X-100 in PBS. The immunostained coverslips were mounted on slides in ProLong Diamond Antifade Mountant with DAPI (ThermoFisher Scientific). Images were acquired with a three-dimensional Zeiss LSM 780 microscopy system (Carl Zeiss) using a 63× oil objective with a numerical aperture of 1.4. Images were collected with a pinhole at 0.7 μm diameter (1 Airy unit), averaged four times, and processed with ZEN 2011 software (Carl Zeiss).

### Crude subcellular fractionation

Fractionation was performed according to the protocol described by Holden and Horton (51) with slight modifications. Transfected cells grown in 100-mm plates were labeled for 30 min with ^35^S and subsequently washed with PBS and detached with 1 ml trypsin. Trypsin was quenched by addition of 2 ml ice-cold complete DMEM media with 10% FBS. Cells were centrifuged for 1 min at 4 °C and 1,000 rpm. Supernatant was aspirated and cells were washed with ice-cold PBS. Collected cells were resuspended in 5 ml ice-cold lysis buffer 1 [150 mM NaCl, 50 mM HEPES, pH 7.4, 25 μg/ml digitonin (Sigma-Aldrich)], incubated for 10 min at 4 °C, and centrifuged for 10 min at 10,000 rpm (step 1). Supernatant was mixed with 500 μl TritonX 114, kept for 1 h on ice, heated for 3 min at 37 °C, and centrifuged for 3 min at 3,000 g at room temperature. The upper aqueous phase contained the purified cytosolic fraction. The pellet from step 1 was washed in cold PBS, resuspended in 5 ml buffer 2 (150 mM NaCl, 50 mM HEPES, pH 7.4, 1% NP40), incubated for 30 min on ice, and centrifuged for 10 min at 2,000 g. The resulting supernatant contained the microsomal fraction. The pellet was resuspended in 2 ml buffer 3 [150 mM NaCl, 50 mM HEPES, pH 7.4, 0.1% SDS, 0.5% sodium deoxycholate, 1 U/ml benzonase (Sigma-Aldrich)], kept for 30 min on ice, and centrifuged for 1 min at 6,000 rpm. The supernatant from this step contained the nuclear fraction. All fractions were examined on Western blots using polyclonal antibodies against individual organelle markers: for the cytosolic fraction, anti-α tubulin antibody (Sigma-Aldrich, #SAB3501071); for the microsomal fraction, anti-PDI antibody (Sigma-Aldrich, #P7496); and for the nuclear fraction, anti-histone H3 antibody (Millipore, #07-690). HBV proteins were immunoprecipitated as described above.

### Co-immunoprecipitation and pull down experiments

Cells co-transfected with either HBe-HA or HBe3CA-HA constructs and DNA encoding individual TRAP c-myc tagged subunits were lysed 48 h post transfection for 1 h in Co-IP buffer (50 mM HEPES, 100 mM NaCl, 10% glycerol, 0.5% DOC, pH 7.9) supplemented with Complete™ EDTA free protease inhibitor cocktail (Roche). Lysates were cleared by centrifugation at 15,000g for 10 min and subjected to immunoprecipitation using anti-HA magnetic beads for 2 h at 4 °C. Collected beads were washed by Co-IP buffer without DOC. Samples were boiled in 2x sample buffer for 5 min to elute the proteins and analyzed by SDS-PAGE followed by Western blot using anti c-myc antibody (Sigma-Aldrich, #C3956).

### Liquid chromatography-tandem mass spectrometry analysis (LC-MS/MS)

Detailed sample preparation has been described previously (52). Briefly, HBe-HA producing cells were treated for 4 h with Brefeldin A (5μg/mL, Sigma-Aldrich) and harvested in lysis buffer containing 50 mM Hepes, 100 mM NaCl, 10% glycerol, 0.5% Nonidet P40, pH 7.9. HA-tagged proteins were immunoprecipitated using anti-HA beads, washed, eluted by HA peptide (0.5 mg/mL), and fragmented using chymotrypsin. The resulting peptides were separated on an UltiMate 3000 RSLCnano system (Thermo Fisher Scientific) coupled to a Mass Spectrometer Orbitrap Fusion Lumos (Thermo Fisher Scientific). The peptides were trapped and desalted with 2% acetonitrile in 0.1% formic acid at a flow rate of 30 μl/min on an Acclaim PepMap100 column [5 μm, 5 mm by 300-μm internal diameter (ID); Thermo Fisher Scientific]. Eluted peptides were separated using an Acclaim PepMap100 analytical column (2 μm, 50-cm by 75-μm ID; ThermoFisher Scientific). The 125-min elution gradient at a constant flow rate of 300 nl/min was set to 5% phase B (0.1% formic acid in 99.9% acetonitrile) and 95% phase A (0.1% formic acid) for the first 1 min. Then, the content of acetonitrile was increased gradually. The orbitrap mass range was set from m/z 350 to 2000 in MS mode, and the instrument acquired fragmentation spectra for ions of m/z 100 to 2000. A Proteome Discoverer 2.4 (ThermoFisher Scientific) was used for peptide and protein identification using Sequest and Amanda as search engines and databases of sequences of HA-tagged HBe or HBe3CA, Swiss-Prot human proteins (downloaded on 15 February 2016), and common contaminants. The data were also searched with MaxQuant (version 1.6.3.4, Max-Planck-Institute of Biochemistry, Planegg, Germany) and the same set of protein databases to obtain peptide and protein intensities applied at the label-free quantification (LFQ) step. Perseus software (version 1.650, Max-Planck-Institute of Biochemistry, Planegg, Germany) was used to perform LFQ comparison of three biological replicates of HA-tagged HBe (or HBe3CA) cells and three biological replicates of cells transfected with empty vector. The data were processed to compare the abundance of individual proteins by statistical tests in form of Student’s *t*-test and resulted in Volcano plot comparing the statistical significance and proteins abundance difference (Fold change).

### HBV particle purification

HBV virions were produced in HepG2-NTCP cells transfected with wt and C3A rcccDNA, as previously described (54). Briefly, rcccDNA for *wt* and mutant precore were transfected in triplicate into HepG2-NTCP cells using Lipofectamine 3000 (ThermoFisher Scientific). The culture medium of transfected cells was collected every three to four days for duration of 30 days. HBV particles were precipitated from clarified cell supernatants by overnight incubation in 6% polyethylene glycol 8000 (PEG 8000) and were then concentrated by centrifugation at 4 °C for 90 min at 14,000 x g. The precipitated virions were re-suspended in complete DMEM supplemented with 10% FBS. HBV titers were determined by quantitative PCR (qPCR) using primers specific for HBV DNA: HBV-F, 5’-AGAGGACTCTTGGACTCTCTGC-3’; HBV-R, 5’-CTCCCAGTCTTTAAACAAACAGTC-3’; and the probe pHBV 5’-[FAM]TCAACGACCGACCTT[BHQ1]-3’. The qPCR reactions were performed with gb Elite PCR Master Mix (Generi Biotech) and TaqMan probe.

### HBV infection and analysis of HBeAg and HBsAg by ELISA

HepG2-NTCP cells were infected with *wt* and C3A mutant HBV in a 12-well plate format. The MOI was 2000 viral genome equivalents per cell. Infection was performed overnight in the presence of 4% PEG8000, 2.5% DMSO, and 3% FBS. The following day, the cells were washed three times with PBS and supplemented with fresh DMEM containing 2.5% DMSO and 3% FBS. The progress of HBV replication was checked by evaluating the titers of HBe and HBs antigens in culture supernatants using a commercial ELISA kit (Bioneovan). Day 0 was defined as the time point after the viral inoculum was washed away and the fresh medium was added to cells.

## Data availability

The mass spectrometry proteomics data have been deposited to the ProteomeXchange Consortium via the PRIDE (53) partner repository with the dataset identifier PXD025430 and 10.6019/PXD025430.

**Password:** hIAHH1G3

## Supporting information

This article contains supporting information

## Acknowledgement

We thank Romana Cubínková for excellent technical support.

## Author contributions

Conceptualization, H.Z. and I.P.; Investigation, H.Z., A.Z., J.H., and B.L; Formal analysis, H.Z., A.Z., B.L., M.H., A.K., and J.W.; Writing – original draft, H.Z., A.Z, and B.L.; Writing – review & editing, J.W. and I.P.; Funding acquisition, J.W. and I.P.

## Abbreviations

HBV: hepatitis B virus
sp: signal peptide
ER: endoplasmic reticulum
HBc: hepatitis B core antigen
HBe: hepatitis B precore antigen
λ-PP: λ-protein phosphatase
DTT: dithiothreitol
TRAP: translocon-associated protein complex

## References

1. Ou JH, Laub O, Rutter WJ. Hepatitis B virus gene function: the precore region targets the core antigen to cellular membranes and causes the secretion of the e antigen. Proc Natl Acad Sci U S A. 1986 Mar;83(6):1578–82.

2. DiMattia MA, Watts NR, Stahl SJ, Grimes JM, Steven AC, Stuart DI, et al. Antigenic switching of hepatitis B virus by alternative dimerization of the capsid protein. Structure. 2013 Jan 8;21(1):133–42.

3. Nassal M, Rieger A. An intramolecular disulfide bridge between Cys-7 and Cys61 determines the structure of the secretory core gene product (e antigen) of hepatitis B virus. J Virol. 1993 Jul;67(7):4307–15.

4. Selzer L, Katen SP, Zlotnick A. The hepatitis B virus core protein intradimer interface modulates capsid assembly and stability. Biochemistry [Internet]. 2014 Sep 2;53(34):5496–504.

5. Jean-Jean O, Salhi S, Carlier D, Elie C, De Recondo AM, Rossignol JM. Biosynthesis of hepatitis B virus e antigen: directed mutagenesis of the putative aspartyl protease site. J Virol. 1989 Dec;63(12):5497–500.

6. Standring DN, Ou JH, Masiarz FR, Rutter WJ. A signal peptide encoded within the precore region of hepatitis B virus directs the secretion of a heterogeneous population of e antigens in Xenopus oocytes. Proc Natl Acad Sci U S A. 1988 Nov;85(22):8405–9.

7. Ito K, Kim K-H, Lok AS-F, Tong S. Characterization of genotype-specific carboxyl-terminal cleavage sites of hepatitis B virus e antigen precursor and identification of furin as the candidate enzyme. J Virol. 2009 Apr;83(8):3507–17.

8. Messageot F, Salhi S, Eon P, Rossignol J. Proteolytic Processing of the Hepatitis B Virus e Antigen Precursor. J Biol Chem. 2003 Jan;278(2):891–5.

9. Chen MT, Billaud J-N, Sällberg M, Guidotti LG, Chisari F V, Jones J, et al. A function of the hepatitis B virus precore protein is to regulate the immune response to the core antigen. Proc Natl Acad Sci U S A. 2004 Oct 12;101(41): 14913–8.

10. Riordan SM, Skinner N, Kurtovic J, Locarnini S, Visvanathan K. Reduced expression of toll-like receptor 2 on peripheral monocytes in patients with chronic hepatitis B. Clin Vaccine Immunol. 2006 Aug;13(8):972–4.

11. Visvanathan K, Skinner NA, Thompson AJ V, Riordan SM, Sozzi V, Edwards R, et al. Regulation of Toll-like receptor-2 expression in chronic hepatitis B by the precore protein. Hepatology. 2007 Jan;45(1):102–10.

12. Lang T, Lo C, Skinner N, Locarnini S, Visvanathan K, Mansell A. The hepatitis B e antigen (HBeAg) targets and suppresses activation of the toll-like receptor signaling pathway. J Hepatol. 2011 Oct;55(4):762–9.

13. Duriez M, Rossigno JM, Sitterlin D. The hepatitis B virus precore protein is retrotransported from endoplasmic reticulum (ER) to cytosol through the ER-associated degradation pathway. J Biol Chem. 2008;283(47):32352–60.

14. Ou JH, Yeh CT, Yen TS. Transport of hepatitis B virus precore protein into the nucleus after cleavage of its signal peptide. J Virol. 1989 Dec;63(12):5238–43.

15. Garcia PD, Ou JH, Rutter WJ, Walter P. Targeting of the hepatitis B virus precore protein to the endoplasmic reticulum membrane: after signal peptide cleavage translocation can be aborted and the product released into the cytoplasm. J Cell Biol. 1988 Apr;106(4):1093–104.

16. Scaglioni PP, Melegari M, Wands JR. Biologic properties of hepatitis B viral genomes with mutations in the precore promoter and precore open reading frame. Virology. 1997;233(2):374–81.

17. Locarnini S, Shaw T, Dean J, Colledge D, Thompson A, Li K, et al. Cellular response to conditional expression of the hepatitis B virus precore and core proteins in cultured hepatoma (Huh-7) cells. J Clin Virol. 2005 Feb;32(2):113–21.

18. Inoue J, Krueger EW, Chen J, Cao H, Ninomiya M, McNiven MA. HBV secretion is regulated through the activation of endocytic and autophagic compartments mediated by Rab7 stimulation. J Cell Sci. 2015 May 1;128(9):1696–706.

19. Liu D, Cui L, Wang Y, Yang G, He J, Hao R, et al. Hepatitis B e antigen and its precursors promote the progress of hepatocellular carcinoma by interacting with NUMB and decreasing p53 activity. Hepatology. 2016 Aug;64(2):390–404.

20. Mitra B, Wang J, Kim ES, Mao R, Dong M, Liu Y, et al. Hepatitis B Virus Precore Protein p22 Inhibits Alpha Interferon Signaling by Blocking STAT Nuclear Translocation. J Virol. 2019 Apr;93(13):e00196–19..

21. Lang S, Pfeffer S, Lee PH, Cavalié A, Helms V, Förster F, et al. An update on Sec 61 channel functions, mechanisms, and related diseases. Front Physiol. 2017 Nov;8:1–22.

22. Alder NN, Shen Y, Brodsky JL, Hendershot LM, Johnson AE. The molecular mechanisms underlying BiP-mediated gating of the Sec61 translocon of the endoplasmic reticulum. J Cell Biol. 2005 Jan;168(3):389–99.

23. Hamman BD, Hendershot LM, Johnson AE. BiP maintains the permeability barrier of the ER membrane by sealing the lumenal end of the translocon pore before and early in translocation. Cell. 1998 Mar;92(6):747–58.

24. Haβdenteufel S, Johnson N, Paton AW, Paton JC, High S, Zimmermann R. Chaperone-Mediated Sec61 Channel Gating during ER Import of Small Precursor Proteins Overcomes Sec61 Inhibitor-Reinforced Energy Barrier. Cell Rep. 2018 May;23(5):1373–86.

25. Pfeffer S, Burbaum L, Unverdorben P, Pech M, Chen Y, Zimmermann R, et al. Structure of the native Sec61 protein-conducting channel. Nat Commun. 2015 Sep 28;6:8403.

26. Hartmann E, Görlich D, Kostka S, Otto A, Kraft R, Knespel S, et al. A tetrameric complex of membrane proteins in the endoplasmic reticulum. Eur J Biochem. 1993;214(2):375–81.

27. Nguyen D, Stutz R, Schorr S, Lang S, Pfeffer S, Freeze HH, et al. Proteomics reveals signal peptide features determining the client specificity in human TRAP-dependent ER protein import. Nat Commun. 2018 Dec;9(1).

28. Fons RD, Bogert BA, Hegde RS. Substrate-specific function of the translocon-associated protein complex during translocation across the ER membrane. J Cell Biol. 2003 Feb;160(4):529–39.

29. Yeh CT, Ou JH. Phosphorylation of hepatitis B virus precore and core proteins. J Virol. 1991 May;65(5):2327–31.

30. Lainé S, Thouard A, Derancourt J, Kress M, Sitterlin D, Rossignol J-M. In Vitro and In Vivo Interactions between the Hepatitis B Virus Protein P22 and the Cellular Protein gC1qR. J Virol. 2003 Dec 1;77(23):12875–80.

31. Heger-Stevic J, Zimmermann P, Lecoq L, Böttcher B, Nassal M. Hepatitis B virus core protein phosphorylation: Identification of the SRPK1 target sites and impact of their occupancy on RNA binding and capsid structure. PLoS Pathog. 2018 Dec 1;14(12).

32. Ludgate L, Adams C, Hu J. Phosphorylation State-Dependent Interactions of Hepadnavirus Core Protein with Host Factors. Ryu W-S, editor. PLoS One. 2011 Dec 22;6(12):e29566.

33. Selzer L, Kant R, Wang JC-Y, Bothner B, Zlotnick A. Hepatitis B Virus Core Protein Phosphorylation Sites Affect Capsid Stability and Transient Exposure of the C-terminal Domain. J Biol Chem. 2015 Nov 20;290(47):28584–93.

34. Wittkop L, Schwarz A, Cassany A, Grün-Bernhard S, Delaleau M, Rabe B, et al. Inhibition of protein kinase C phosphorylation of hepatitis B virus capsids inhibits virion formation and causes intracellular capsid accumulation. Cell Microbiol. 2010 Jan 26;12(7):962–75.

35. Liao W, Ou JH. Phosphorylation and nuclear localization of the hepatitis B virus core protein: significance of serine in the three repeated SPRRR motifs. J Virol. 1995 Feb;69(2):1025–9.

36. Rabe B, Vlachou A, Pante N, Helenius A, Kann M. Nuclear import of hepatitis B virus capsids and release of the viral genome. Proc Natl Acad Sci. 2003 Aug;100(17):9849–54.

37. Lubyova B, Hodek J, Zabransky A, Prouzova H, Hubalek M, Hirsch I, et al. PRMT5: A novel regulator of Hepatitis B virus replication and an arginine methylase of HBV core. PLoS One. 2017 Oct;12(10):e0186982.

38. Li HC, Huang EY, Su PY, Wu SY, Yang CC, Lin YS, et al. Nuclear export and import of human hepatitis B virus capsid protein and particles. PLoS Pathog. 2010 Oct;6(10).

39. Yeh CT, Hong LH, Ou JH, Chu CM, Liaw YF. Characterization of nuclear localization of a hepatitis B virus precore protein derivative P22. Arch Virol. 1996;141(3–4):425–38.

40. Wasenauer G, Köck J, Schlicht HJ. A cysteine and a hydrophobic sequence in the noncleaved portion of the pre-C leader peptide determine the biophysical properties of the secretory core protein (HBe protein) of human hepatitis B virus. J Virol. 1992 Sep;66(9):5338–46.

41. Shaffer KL, Sharma A, Snapp EL, Hegde RS. Regulation of protein compartmentalization expands the diversity of protein function. Dev Cell. 2005 Oct;9(4):545–54.

42. Levine CG, Mitra D, Sharma A, Smith CL, Hegde RS. The efficiency of protein compartmentalization into the secretory pathway. Mol Biol Cell. 2005 Jan;16(1):279–91.

43. Rapoport TA. Protein translocation across the eukaryotic endoplasmic reticulum and bacterial plasma membranes. Nature. 2007 Nov 29;450(7170):663–9.

44. Bonardi F, Halza E, Walko M, Du Plessis F, Nouwen N, Feringa BL, et al. Probing the SecYEG translocation pore size with preproteins conjugated with sizable rigid spherical molecules. Proc Natl Acad Sci U S A. 2011 May;108(19):7775–80.

45. Arkowitz RA, Joly JC, Wickner W. Translocation can drive the unfolding of a preprotein domain. EMBO J. 1993 Jan;12(1):243–53.

46. Hegde RS, Kang SW. The concept of translocational regulation. Vol. 182, Journal of Cell Biology. The Rockefeller University Press; 2008. p. 225–32.

47. Yang SQ, Walter M, Standring DN. Hepatitis B virus p25 precore protein accumulates in Xenopus oocytes as an untranslocated phosphoprotein with an uncleaved signal peptide. J Virol. 1992 Jan;66(1):37–45.

48. Kriegler T, Lang S, Notari L, Hessa T. Prion Protein Translocation Mechanism Revealed by Pulling Force Studies. J Mol Biol. 2020 Jul;432(16):4447–65.

49. Kriegler T, Kiburg G, Hessa T. Translocon-associated protein complex (TRAP) is crucial for co-translational translocation of pre-proinsulin. J Mol Biol. 2020 Dec 4;432(24):166694.

50. Guo X, Chen P, Hou X, Xu W, Wang D, Wang TY, et al. The recombined cccDNA produced using minicircle technology mimicked HBV genome in structure and function closely. Sci Rep. 2016 May 13;6:25552.

51. Holden P, Horton WA. Crude subcellular fractionation of cultured mammalian cell lines. BMC Res Notes. 2009 Jan;2:243.

52. Langerová H, Lubyová B, Zábranský A, Hubálek M, Glendová K, Aillot L, et al. Hepatitis B Core Protein Is Post-Translationally Modified through K29-Linked Ubiquitination. Cells. 2020 NOV 26;9(12):2547.

53. Perez-Riverol Y, Csordas A, Bai J, Bernal-Llinares M, Hewapathirana S, Kundu DJ, et al. The PRIDE database and related tools and resources in 2019: Improving support for quantification data. Nucleic Acids Res. 2019 Jan 8;47(D1):D442–50.

54. Ni Y, Sonnabend J, Seitz S, Urban S. The Pre-S2 Domain of the Hepatitis B Virus Is Dispensable for Infectivity but Serves a Spacer Function for L-Protein-Connected Virus Assembly. J Virol. 2010 Apr;84(8):3879–88.

